# Platycodin D prevents both lysosome- and TMPRSS2-driven SARS-CoV-2 infection *in vitro* by hindering membrane fusion

**DOI:** 10.1101/2020.12.22.423909

**Authors:** Tai Young Kim, Sangeun Jeon, Youngho Jang, Lizaveta Gotina, Joungha Won, Yeon Ha Ju, Sunpil Kim, Minwoo Wendy Jang, Woojin Won, Mingu Gordon Park, Ae Nim Pae, Sunkyu Han, Seungtaek Kim, C. Justin Lee

**Affiliations:** Center for Cognition and Sociality, Cognitive Glioscience Group, Institute for Basic Science, Daejeon 34126, Korea; Zoonotic Virus Laboratory, Institut Pasteur Korea, Seongnam, Korea; Department of Chemistry, Korea Advanced Institute of Science and Technology (KAIST), Daejeon 34141, Korea; Convergence Research Center for Diagnosis, Treatment and Care System of Dementia, Korea Institute of Science and Technology, Hwarangno 14-gil 5, Seongbuk-gu, Seoul, 02792, Republic of Korea; Division of Bio-Medical Science & Technology, KIST School, Korea University of Science and Technology (UST), Daejeon, Korea; Department of Biological Sciences, Korea Advanced Institute of Science and Technology (KAIST), Daejeon 34141, Korea; IBS School, University of Science and Technology, Daejeon, Republic of Korea; Neuroscience Program, University of Science and Technology, Daejeon, Republic of Korea; KU-KIST Graduate School of Converging Science and Technology, Korea University, Seoul 02841, Korea

**Keywords:** *Platycodon grandiflorum*, Platycodin D, natural product, SARS-CoV-2, COVID-19, lysosome, TMPRSS2, cholesterol distribution, membrane fusion

## Abstract

An ongoing pandemic of coronavirus disease 2019 (COVID-19) is now the greatest threat to the global public health. Herbal medicines and their derived natural products have drawn much attention to treat COVID-19, but there has been no natural product showing inhibitory activity against SARS-CoV-2 infection with detailed mechanism. Here, we show that platycodin D (PD), a triterpenoid saponin abundant in *Platycodon grandiflorum* (PG), a dietary and medicinal herb commonly used in East Asia, effectively blocks the two main SARS-CoV-2 infection-routes via lysosome- and transmembrane protease, serine 2 (TMPRSS2)-driven entry. Mechanistically, PD prevents host-entry of SARS-CoV-2 by redistributing membrane cholesterol to prevent membrane fusion, which can be reinstated by treatment with a PD-encapsulating agent. Furthermore, the inhibitory effects of PD are recapitulated by a pharmacological inhibition or gene-silencing of *NPC1*, which is mutated in Niemann-Pick type C (NPC) patients displaying disrupted membrane cholesterol. Finally, readily available local foods or herbal medicines containing PG root show the similar inhibitory effects against SARS-CoV-2 infection. Our study proposes that PD is a potent natural product for preventing or treating COVID-19 and that a brief disruption of membrane cholesterol can be a novel therapeutic approach against SARS-CoV-2 infection.

## Introduction

SARS-CoV-2 expresses spike glycoprotein (S) on their surface and uses it to bind to the host receptor, angiotensin-converting enzyme 2 (ACE2) (Hoffmann et al., 2020a; Li et al., 2003). SARS-CoV-2 is known to enter host cells during infection via two pathways: 1) the endocytic pathway followed by a cathepsin B/L-mediated cleavage of S protein in lysosomes, and 2) a direct fusion of the virus envelope with the host plasma membrane after a TMPRSS2-mediated cleavage of S protein (Cai et al., 2020; Hoffmann et al., 2020a). The cleavage of S protein by cathepsin B/L or TMPRSS2 or both is the critical step to release the viral RNA into the cytosol of host cells. Therefore, the abundance and availability of the two host proteases, cathepsin B/L and TMPRSS2, and the host receptor, ACE2, are the most critical determining factor for the host susceptibility to COVID-19. For example, chloroquine, a well-known anti-malaria drug, was initially suggested to be a potent anti-SARS-CoV-2 drug due to its ability to block the lysosomal cathepsin-dependent virus-entry by elevating the lysosomal pH (Liu et al., 2020; Wang et al., 2020a). Unfortunately, the most of clinical trials with chloroquine and hydroxychloroquine failed to show beneficial effects in COVID-19 patients (Pathak et al., 2020). The failure of chloroquine was attributed to its inability to block the TMPRSS2-mediated SARS-CoV-2-entry (Hoffmann et al., 2020b). Subsequently, TMPRSS2 inhibitors such as camostat and nafamostat emerged as the next generation blockers of SARS-CoV-2-entry. However, they also fell short due to their inability to block SARS-CoV-2-entry into certain cell-types that express only ACE2 without TMPRSS2 (Ko et al., 2020; Yamamoto et al., 2020). Therefore, discovery and development of drugs that block both lysosome- and TMPRSS2-driven SARS-CoV-2-entry are desperately needed.

Herbal medicines and their derived natural products have drawn much attention to treat COVID-19 because they have been shown to possess antiviral properties against a broad range of pathogenic viruses including influenza, HIV, SARS-CoV, and MERS-CoV (Huang et al., 2020). To date, numerous natural products have been proposed to inhibit one of the essential components of SARS-CoV-2, including main protease (Mpro), RNA-dependent RNA polymerase (RdRp), ACE2, and TMPRSS2, although most of them are based on *in silico* screening through molecular docking approaches (Joshi et al., 2020; Khan et al., 2020; Vivek-Ananth et al., 2020). Only very few of them have been verified for their inhibitory activity (Senthil Kumar et al., 2020), and none of them have been reported with detailed molecular mechanism of action. Among thousands of Korean traditional herbal medicines, we have focused on the root of *Platycodon grandiflorum* (PG) (**Figure 1A**), which has been described in the Dongui Bogam, the most famous 17^th^ century Korean medical textbook (Song et al., 2016). The Dongui Bogam reports that PG can be used to treat patients with disorders in respiratory tracts and the lung, the major target sites of SARS-CoV-2 (Ziegler et al., 2020). Even in the present day, PG is widely used in East Asia including Korea, China, and Japan to treat several respiratory diseases such as asthma, airway inflammation, and sore throats (Choi et al., 2009; Ishimaru et al., 2013; Shin et al., 2002). Several studies have demonstrated that the root of PG is enriched with platycodin D (PD) (**Figure 1B**), a glycosylated triterpenoid saponin (colored in blue; **Figure 1B**), which is the major active natural product in mediating those biological activities (Nyakudya et al., 2014; Zhang et al., 2015). Recently, PD has been reported to exhibit a potent antiviral activity against Type 2 porcine reproductive and respiratory syndrome virus (PRRSV) and Hepatitis C virus (HCV) (Kim et al., 2013; Zhang et al., 2018). However, its inhibitory activity against coronaviruses and its mechanism of action have not been explored. In this study, we set out to investigate whether PG and PD show an anti-SARS-CoV-2 activity by blocking both lysosome- and TMPRSS2-driven SARS-CoV-2-entry.

**Figure 1.**
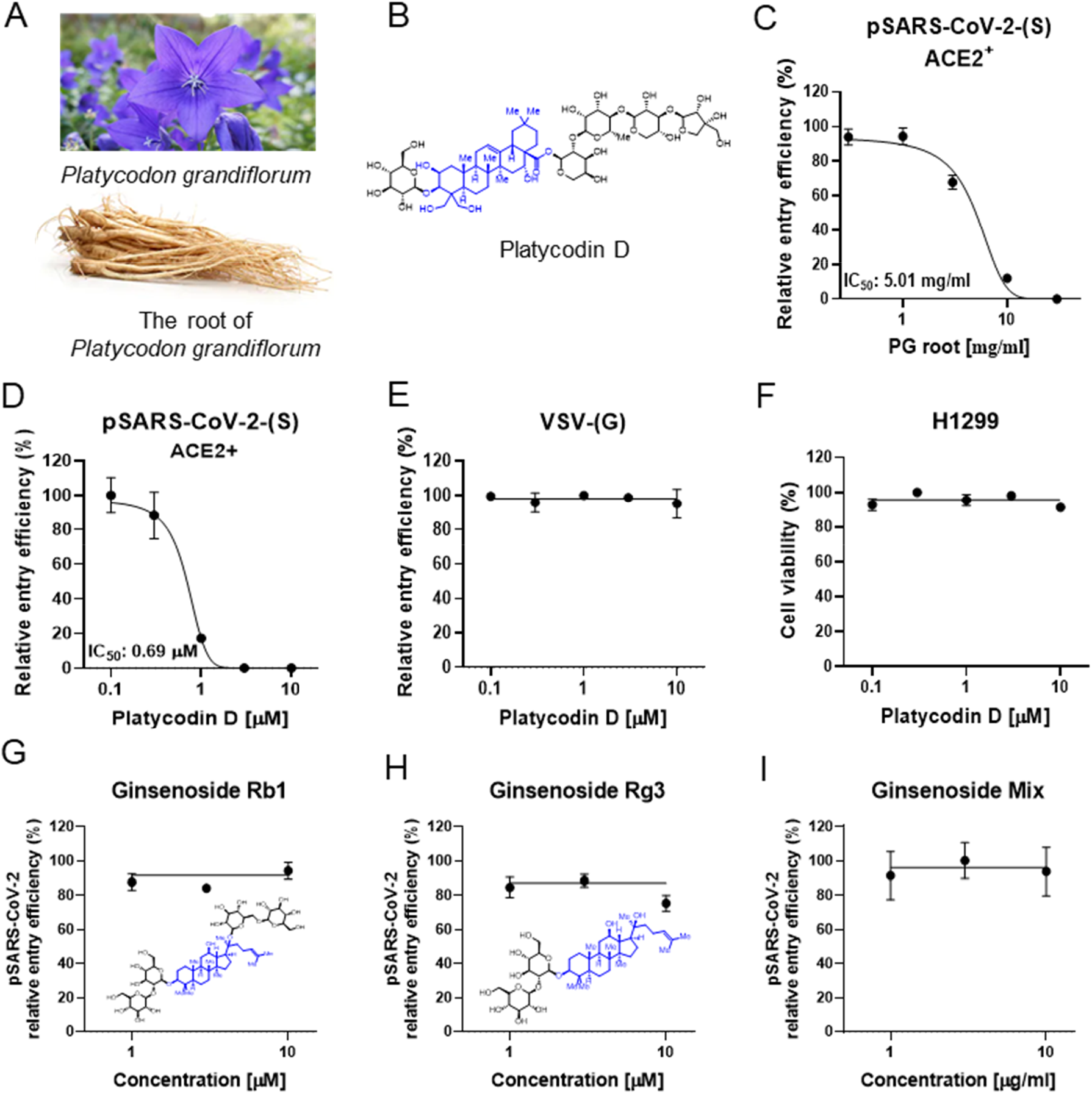
PD, a saponin present only in *Platycodon grandiflorum*, blocks SARS-CoV-2 entry into human cells. **(A)** *Platycodon grandiflorum* (balloon flower) and the root of *Platycodon grandiflorum*. **(B)** Chemical structure of Platycodin D with triterpenoid saponin in blue. **(C-F)** ACE2^+^ H1299 cells pre-treated for 1 h with the indicated concentrations of water extract of PG root or PD and were transduced with pSARS-CoV-2-(S) (C and D) or VSV-(G) (E). Cytotoxicity was determined using WST-8 cell viability assay (F). **(G-I)** pSARS-CoV-2 entry assay with ginsenoside Rb1 (1, 3, 10 μM) (G), R3 (1, 3, 10 μM) (H) or ginsenoside mixture (1, 3, 10 μg/ml) (I). Inset shows the chemical structure of Rb1 and Rg3 (G, H). **(C-E, G-I)**. After culture for 24 h, transduction efficiency was quantified by measuring the activity of firefly luciferase in cell lysates. Data are representative of three independent experiments with triplicate samples. Error bars indicate SEM.

## Results

### PD, a triterpenoid saponin present in Platycodon grandifloras, has a specific inhibitory activity against SARS-CoV-2 entry

To test for an anti-SARS-CoV-2 activity, we first developed a SARS-CoV-2 pseudovirus (pSARS-CoV-2), which carried the full-length S protein of SARS-CoV-2 on the HIV-based lentiviral particles and the luciferase gene as a reporter (Crawford et al., 2020), to mimic the S protein of the native SARS-CoV-2 virus and retain its ability to bind to the host cell surface receptors for viral infection. We then examined whether PG and PD can prevent pSARS-CoV-2-entry into H1299, a human lung cell line known to be susceptible to coronavirus infection (Tay et al., 2012). Luciferase activity assay after infection with pSARS-CoV-2 revealed that virus-entry into H1299 required overexpression of ACE2 (ACE2^+^) (**Figure S1**). In ACE2^+^, we found that 1-hour treatment with PG and PD effectively reduced pSARS-CoV-2-entry in a dose-dependent manner with a half-maximal inhibitory concentration (IC_50_) of 5.01 mg/ml (**Figure 1C**) and 0.69μM (**Figure 1D**), respectively. In contrast, PD did not block the entry of the control lentiviral particles driven by the glycoprotein (G proteins) of the vesicular stomatitis virus (VSV) (**Figure 1E**). PD showed no cytotoxic effect on H1299 *per se* at the tested concentrations (**Figure 1F**). These results indicate that the inhibitory effect of PD on virus-entry requires both S protein of SARS-CoV-2 and ACE2 in the host cells. To test the specificity of PD among other saponins, we compared it with ginsenosides, which are a group of saponins from *Panax ginseng*, also known as Korean ginseng. Ginsenosides are known to exhibit antiviral activity against multiple types of virus such as rhinovirus, influenza virus, HIV, hepatitis virus, and herpesvirus (Im et al., 2016). In the pSARS-CoV-2 entry assay, none of the ginsenosides we tested including Rb1, Rg3, and ginsenoside mixture, prevented pSARS-CoV-2 from entering ACE2^+^ (**Figure 1G-I**), suggesting that PD, a triterpenoid saponin only present in *Platycodon grandifloras*, possesses a specific inhibitory activity against infection of SARS-CoV-2.

### Herbal medicine and foods containing PG inhibit both lysosome and TMPRSS2-mediated SARS-CoV-2 entry pathways through the action of PD

To determine which one of the two entry pathways is the target of PD, we prepared additional cell lines, H1299 and HEK293T, overexpressing both ACE2 and TMPRSS2 (ACE2/TMPRSS2^+^) and compared the inhibitory effect of various drugs with ACE2^+^. We found that E64d and chloroquine, inhibitors of lysosomal cathepsins, effectively blocked pSARS-CoV-2-entry only in ACE2^+^, but not in ACE2/TMPRSS2^+^, whereas camostat and nafamostat, inhibitors of TMPRSS2, effectively blocked pSARS-CoV-2-entry in ACE2/TMPRSS2^+^, but not in ACE2^+^ (**Figure 2A, B**). In contrast, 5 μM PD completely inhibited pSARS-CoV-2 entry in both ACE2/TMPRSS2^+^ and ACE2^+^. (**Figure 2A, B**). The actual potency of PD for inhibition of pSARS-CoV-2-entry in terms of IC_50_’s in ACE2/TMPRSS2^+^ was determined to be 0.72 μM (**Figure 2C**), which was almost identical to that in ACE2^+^ (**Figure 1D**). These results suggest that PD targets any one of the events that is common to both entry pathways.

**Figure 2.**
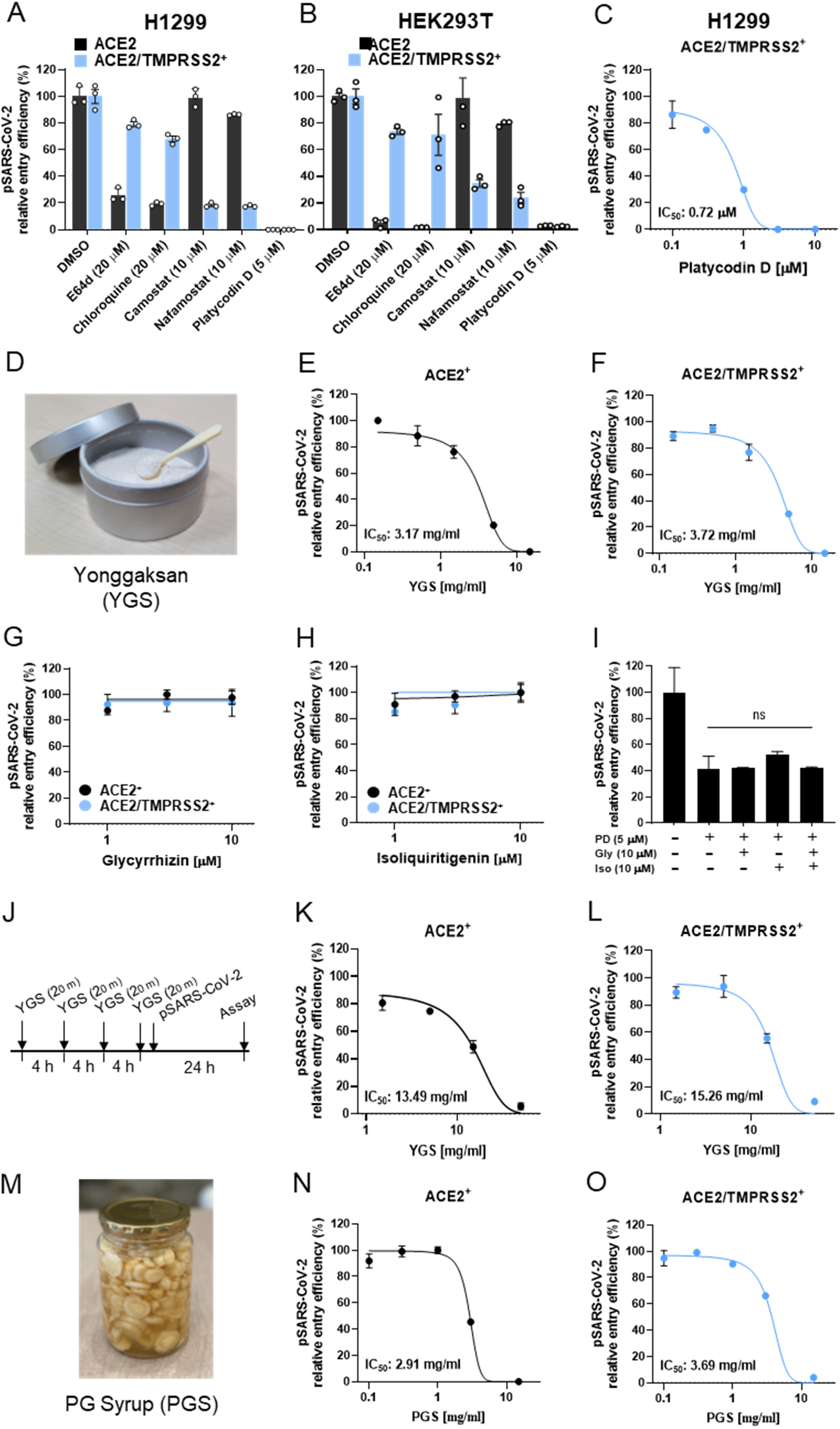
PD and PG root-containing herbal medicine and foods blocks two main SARS-CoV-2 entry pathways. **(A, B)** To analyze whether PD inhibits lysosome- or TMPRSS2-dependent cell entry, H1299 (A) and HEK293T (B) cells that express ACE2 alone (ACE2^+^) or in combination with TMPRSS2 (ACE2/TMPRSS2^+^) were pre-treated with 5 μM PD for 1 h prior to transduction with pSARS-CoV-2 in the presence of PD. Lysosomal protease inhibitors E64d and chloroquine, and TMPRSS2 inhibitors, camostat and nafamostat, were used to verify pSARS-CoV-2-entry pathways in these cells. **(C)** ACE2/TMPRSS2^+^ H1299 cells were pretreated for 1 h with the indicated concentrations of PD and transduced with pSARS-CoV-2 in the presence of PD. **(D)** Yonggaksan (YGS) is composed of a group of herbal powders including PG root. **(E-I)** ACE2^+^ and ACE2/TMPRSS2^+^ were treated with serial three-fold dilutions of YGS stock solution (15 mg/ml) (E, F), 1, 3, 10 μM of glycyrrhizin and isoliquiritigenin (G, H), or 1 μM PD in the absence or presence of 10 μM of glycyrrhizin, isoliquiritigenin, or glycyrrhizin plus isoliquiritigenin (I) for 1 h prior to transduction with pSARS-CoV-2 in the presence of each drugs. **(J)** Experimental timeline for YGS pretreatment before transduction with pSARS-CoV-2. **(K, L)** ACE2^+^ and ACE2/TMPRSS2^+^ were pretreated with serial three-fold dilutions of YGS with a starting concentration of 100 mg/ml for 20 min, four times with 4 h interval, and then transduced with pSARS-CoV-2 without YGS. (**M**) PG syrup, mainly made of PG root and frequently used for respiratory disease. **(N, O)** ACE2^+^ and ACE2/TMPRSS2^+^were pretreated with serial three-fold dilutions of PG syrup stock solution containing 15 mg/ml of the PG root for 1 h prior to transduction with pSARS-CoV-2 in the presence of the syrup. **(A-C, E-I, K, L, N, O)** After culture for 24 h, transduction efficiency was quantified by measuring the activity of firefly luciferase in cell lysates. Data are representative of two or three independent experiments with triplicate samples. Error bars indicate SEM. P values were determined by unpaired, two-tailed Student’s t-test. NS, not significant.

Yonggaksan (YGS, **Figure 2D**), whose main ingredient is an extract powder from the PG root, has been available as an over-the-counter medicine and used for treatment of phlegm, cough, and sore throat for more than 50 years in Korea and 200 years in Japan. Thus, we examined whether YGS can be a potential nonprescription medicine to block SARS-CoV-2 infection. We found that a dilute solution of YGS effectively inhibited pSARS-CoV-2-entry into both ACE2^+^ and ACE2/TMPRSS2^+^ in a dose-dependent manner with similar IC_50_’s of 3.17 mg/ml in ACE2^+^ (**Figure 2E**) and 3.72 mg/ml in ACE2/TMPRSS2^+^ (**Figure 2F**), suggesting a therapeutic potential of YGS for SARS-CoV-2. Another main ingredient of YGS is an extract powder from the liquorice roots, whose active compound is glycyrrhizin with an inhibitory action in replication and penetration of SARS-CoV (Cinatl et al., 2003) and a suggested therapeutic potential for COVID-19 (Luo et al., 2020). Thus, we tested the effect of glycyrrhizin as well as isoliquiritigenin, other major components of liquorice roots, on the entry of pSARS-CoV-2. We found that both compounds exhibited no significant inhibitory activity against pSAR-CoV-2-entry in both ACE2^+^ and ACE2/TMPRSS2^+^ (**Figure 2G, H**). Moreover, mixing the two compounds with 1 μM PD did not either enhance or reduce pSARS-CoV-2-entry in ACE2^+^ (**Figure 2I**). Quantification of components in YGS by high performance liquid chromatography (HPLC) further revealed that PD is the main active compound of YGS to block pSARS-CoV-2-entry (**Figure S2, S3**). The common daily dosage and regimen of YGS are by taking one spoonful (0.3 gram) of YGS powder on the throat surface without water, waiting for 20 ~ 30 minutes, and repeating 3 ~ 6 times a day. To mimic this usage, we treated the cells with YGS four times for 20 min each time at 4-hour interval before adding pSARS-CoV-2 (**Figure 2J**). Under this dosage and regimen, YGS effectively reduced pSARS-CoV-2-entry with similar IC_50_’s of 13.49 mg/ml in ACE2^+^ (**Figure 2K**) and 15.26 mg/ml in ACE2/TMPRSS2^+^ (**Figure 2L**). Taken together, these results suggest that YGS, through the action of PD, can be a high potential, nonprescription, over-the-counter medicine to fight against SARS-CoV-2 infection.

In Korean culture, the root of PG is often served as a side dish in daily meals. For example, PG syrup (PGS) has been considered as a folk remedy for relieving several symptoms of respiratory diseases. PGS (**Figure 2M**) is prepared by chopping PG roots (**Figure 1A**) into small pieces and marinating them in sugar or honey for several months to be mixed with warm water and served as a drink. To test if PGS also shows inhibitory effect on pSARS-CoV-2-entry, we treated ACE2^+^ and ACE2/TMPRSS2^+^ with serial three-fold dilutions of PGS stock solution containing 15 mg/ml of the PG root for 1 h, before infection of pSARS-CoV-2. We found that PGS effectively inhibited pSARS-CoV-2-entry with similar IC_50_’s of 2.91 mg/ml (**Figure 2N**) in ACE2^+^ and 3.69 mg/ml in ACE2/TMPRSS2^+^ (**Figure 2O**). Moreover, HPLC analysis demonstrated that PD is the main active compound in PG root with inhibitory activity against pSARS-CoV-2-entry (**Figure S2, S3**). These results suggest that even dietary foods containing PG roots might also have beneficial effects on COVID-19 patients.

### PD blocks SARS-CoV-2 entry by preventing cholesterol-dependent membrane fusion

To delineate the detailed molecular and cellular mechanism of PD action, we explored two possible events that are common to both entry pathways; 1) the initial binding of S protein to ACE2 at the plasma membrane and 2) the fusion of viral membrane onto host-cell membrane for the entry of viral RNA into the cytosol of host cell. Of the two events, we eliminated the possibility of PD influencing a specific interaction between SARS-CoV-2 S protein and ACE2 by using the GFP-tagged receptor binding domain (RBD) of S protein and FACS analysis (**Figure 3A**). We then investigated the possible mechanism of PD action during the membrane fusion. It has been reported that PD helps lower cholesterol levels in the mouse model of hypercholesterolemia (Zhao et al., 2006) and that depletion of membrane cholesterol content inhibits TMPRSS2-mediated coronavirus-fusion (Wang et al., 2020b; Zang et al., 2020; Zu et al., 2020), raising a possibility that PD directly influences the membrane cholesterol content to disturb membrane fusion. To test the role of cholesterol in SARS-CoV-2 entry, we performed the entry assay with the ACE2^+^ grown in lipoprotein-free culture media and compared to the control media. Lipoprotein-free culture conditions caused a significant reduction of pSARS-CoV-2 entry in ACE2^+^ (**Figure 3B**) and a significant depletion of cholesterol in plasma membrane as well as in other intracellular compartments revealed by staining of ACE2^+^ with filipin-III, a fluorescent dye that binds to free cholesterol (**Figure 3C**). These results support the idea that PD’s inhibitory action on SARS-CoV-2 entry could be due to its effect on the membrane cholesterol.

**Figure 3.**
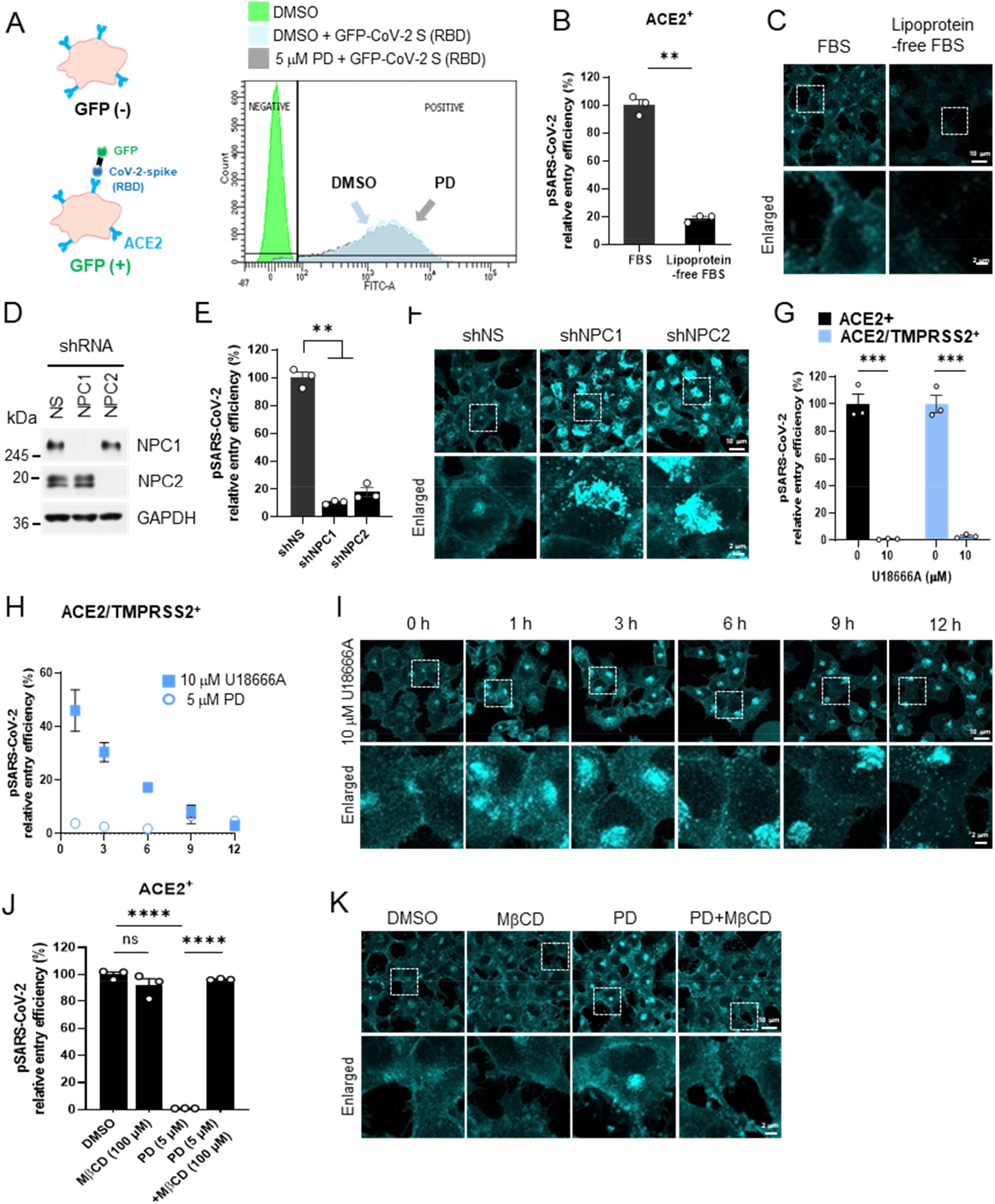
PD blocks SARS-CoV-2-entry by preventing cholesterol-dependent membrane fusion. **(A)** Schematic illustration showing that ACE2^+^ become GFP-positive when GFP-conjugated SARS-CoV-2 spike receptor binding domain (RBD), GFP-CoV-2 S (RBD) is bound to ACE2 on host cells. ACE2^+^pre-treated with DMSO or 5 μM PD for 1 h were incubated with DMSO or GFP-CoV-2 S (RBD), followed by FACS analysis to examine the effects of PD on the SARS-CoV-2 binding to cell surface ACE2. **(B, C)** ACE2^+^ were incubated in culture media containing 10% FBS or 10% lipoprotein-fee FBS for 24 h prior to transduction with pSARS-CoV-2 (A), or staining with filipin-III (B). **(D-F)** ACE2^+^ were infected with lentiviral particles expressing non-specific (NS) shRNA, or shRNA targeting *NPC1* or *NPC2* for 4 days, and then the protein levels of NPC1 and NPC2 were assessed by western blot analysis. GAPDH was used as a loading control (D). The cells were further transduced with pSARS-CoV-2 (E) or stained with filipin-III (F). **(G)** ACE2^+^ and ACE2/TMPRSS2^+^ were pre-treated with 10 μM U18666A for 12 h prior to transduction with pSARS-CoV-2. **(H, I)** ACE2/TMPRSS2^+^pretreated with 10 μM U18666A or 5 μM PD for 1, 3, 6, 9, 12 h, followed by transduction with pSARS-CoV-2 (H) or staining with filipin-III (I). **(J, K)** ACE2^+^ were treated with 5 μM PD in the presence or absence of 100 μM MβCD for 1 h and transduced with pSARS-CoV-2 in the presence of the same concentration of drugs (J) or stained with filipin-III (K). **(B, E, G, H, J)** After culture for 24 h, transduction efficiency was quantified by measuring the activity of firefly luciferase in cell lysates. Data are representative of two or three independent experiments with triplicate samples. Error bars indicate SEM. P values were determined by unpaired, two-tailed Student’s t-test (B, E, G) or one-way analysis of variance (ANOVA) using Tukey’s post hoc test (J). *P < 0.05; **P < 0.01; ***P < 0.001; ****P < 0.0001; NS, not significant. Error bars are not visible where they are within the symbols (H).

As another example of disrupted cholesterol trafficking, we explored the cellular model of Niemann-Pick type C (NPC), which is characterized by a redistribution of free cholesterol from the plasma membrane and an accumulation in the late endosome and lysosome compartment (Carstea et al., 1997). NPC is caused by loss-of-function mutations in either *NPC1* or *NPC2* gene (Sleat et al., 2004). After generating *NPC1* or *NPC2* specific shRNA (**Figure 3D**), we performed pSARS-CoV-2-entry assay and found that gene-silencing of *NPC1* or *NPC2* by each shRNA caused a significant reduction in pSARS-CoV-2 entry in ACE^+^ (**Figure 3E**). Staining with filipin-III showed the expected altered trafficking of membrane cholesterol and accumulated endolysosomal cholesterol (**Figure 3F**). We next utilized U18666A, a well-known inhibitor of NPC1 to pharmacologically disturb the membrane cholesterol. U18666A is known to inhibit NPC1 function by directly binding to sterol-sensing domain of NPC1 protein (Lu et al., 2015). Furthermore, treatment with U18666A has been shown to disrupt the entry of a variety of enveloped viruses, including Dengue virus, Ebola virus, Hepatitis C virus, Influenza A virus, Zika virus, Chikungunya virus, and HIV (Salata et al., 2017), as well as coronaviruses such as SARS-CoV, Middle East respiratory syndrome-related coronavirus (MERS-CoV), and Type I Feline Coronavirus (F-CoV) (Takano et al., 2017; Wrensch et al., 2014). We found that 12-hour treatment with 10 μM U18666A completely blocked pSARS-CoV-2-entry in both ACE2^+^ and ACE2/TMPRSS2^+^ (**Figure 3G**). Intriguingly, the slow time course of blocking effect by U18666A on pSARS-CoV-2-entry in ACE2/TMPRSS2^+^ (**Figure 3H**) coincided very well with the slow time course of the disappearance of membrane cholesterol (**Figure 3I**). This slow time course of U18666A was in marked contrast to the fast-acting effect of PD (**Figure 3H**). These results suggest that disturbing the membrane cholesterol can be an effective target to block SARS-CoV-2-entry.

To test if PD blocks SARS-CoV-2-entry by disrupting the membrane cholesterol, we utilized methyl β-cyclodextrin (MβCD), whose doughnut-shaped structure makes it a natural complexing agent that can encapsulate cholesterol and other large molecules such as hormones, vitamins and natural hydrophobic macro-compounds (Dai et al., 2017). It has been demonstrated that when cells are exposed to high concentration of MβCD (5-10 mM) for more than 2 hours, 80–90% of total cellular cholesterol can be removed (Zidovetzki and Levitan, 2007). However, at much lower concentrations and 1-hour incubation time, MβCD should not encapsulate cholesterol. Indeed, under the condition of 100 μM and 1-hour incubation time, MβCD treatment showed no significant effect on pSARS-CoV-2-entry in ACE2^+^ (**Figure 3J**). Under the same condition, MβCD completely reverted the blocking effect of PD on pSARS-CoV-2-entry to the level of control DMSO condition (**Figure 3J**). PD alone induced a dramatic redistribution of membrane and endolysosomal cholesterol (**Figure 3K**), which was somewhat different from the U18666A’s effect (**Figure 3I**, 12 h). These results indicate that 100 μM MβCD completely sequestered 5 μM PD, without affecting the membrane cholesterol content. Indeed, staining with filipin-III showed no alteration of membrane cholesterol by 100 μM MβCD alone, a significant redistribution of membrane cholesterol by 5 μM PD, and complete reversion by 100 μM MβCD plus 5 μM PD condition (**Figure 3J**).

To understand the detailed molecular mechanisms of how PD prevents pSARS-CoV-2-entry at the level of chemical structure of PD, we utilized the dynamic molecular modelling techniques. Based on the previously reported crystal structure of cholesterol-βCD encapsulation complex (**Figure 4A**) (Christoforides, 2018), we performed a homology-modeling of cholesterol-MβCD and PD-MβCD complexes (**Figure 4D, E**) and calculated the binding energy for each molecular pair in terms of Glide G score and Molecular Mechanics/Generalized Born Surface Area (MM-GBSA) free energy scores (**Figure 4F**). We found that out of 21 hydroxyl groups in βCD (**Figure 4B**), 12 ~ 18 hydroxyl groups could be methylated at different positions, while MβCD with 14 methyl groups showed the greatest encapsulating capacity. Therefore, we chose tetradeca-2,6-O-methyl-β-cyclodextrin (**Figure 4C**) as the representative MβCD to model the cholesterol-MβCD and PD-MβCD complexes. The cholesterol-MβCD encapsulation complex was formed by two MβCD molecules arranged co-axially, forming a head-to-head dimer via intra-molecular hydrogen bonds and one cholesterol molecule fitted snugly in the hydrophobic cavity of MβCD, whose methoxymethyl groups encapsulated cholesterol better in comparison to the cholesterol-βCD complex (**Figure 4D**). In the PD-MβCD encapsulation complex, the hydrophobic core of PD fitted well into the MβCD host dimer, whereas PD’s hydrophilic sugar tail was fully exposed to the solvent (**Figure 4E**). Overall, the molecular modeling of MβCD, PD, and cholesterol revealed that a homodimer of MβCD snugly encapsulated PD and cholesterol with a higher affinity to PD than cholesterol (**Figure 4F**), supporting the possibility that MβCD preferentially sequesters PD over cholesterol. In particular, the predicted orientation of PD revealed that the hydrophobic triterpenoid saponin moiety of PD was completely encapsulated within the homodimeric MβCD capsule, whereas the hydrophilic sugar moiety was extruded through a hole of the capsule (**Figure 4E**). A similar molecular modeling of membrane-inserted PD and cholesterol in an explicit lipid bilayer model also predicted the similar orientation of the partly-embedded PD and fully-embedded cholesterol (**Figure 4G**). In particular, unlike cholesterol, the hydrophilic sugar moiety of PD was sticking out of the lipid bilayer (**Figure 4G**), creating a protrusion on the surface of membrane, which might act as a hindrance to membrane fusion. Furthermore, none of other natural triterpene compounds without any sugar moiety but with similar backbone structure as PD, including echinocystic acid, oleanolic acid, and ursolic acid, showed inhibitory activity against pSARS-CoV-2 entry (**Figure 4H, I**) and altered distribution of membrane cholesterol (**Figure 4J**), implying that the sugar moieties attached to the saponin backbone of PD might be responsible for its action against pSARS-CoV-2-entry via altering membrane cholesterol. These results raise a strong possibility that PD might prevent membrane fusion in a cholesterol-dependent manner.

**Figure 4.**
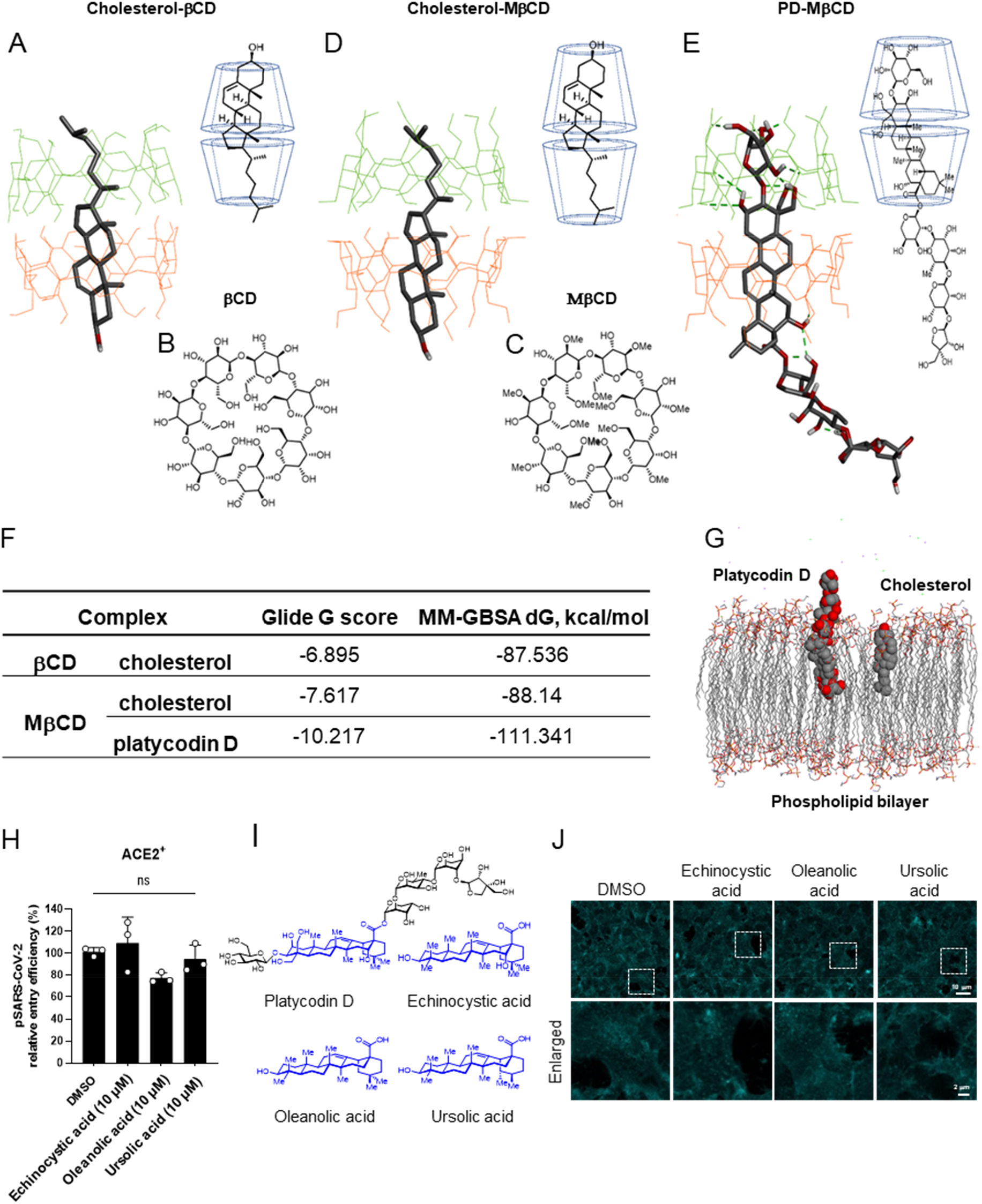
The encapsulation of PD by MβCD and significantly higher affinity of MβCD to PD than to cholesterol. **(A)** Energy minimized cholesterol-βCD crystal structure. **(B, C)** Chemical structure of βCD (B) and tetradeca-2,6-O-methyl-β-cyclodextrin (C). **(D, E)** cholesterol-MβCD inclusion complex, modelled based on the βCD template (D). PD-MβCD inclusion complex (E). Guest ligand is in stick form, whereas host molecules are presented as green and orange lines. Hydrogen bonds (both intramolecular and those between the host and guest molecules) are depicted as green dashed lines. Only polar hydrogens are shown for image clarity **(F)** Glide scores and binding affinity calculations for βCD inclusion complexes (**G**) Model of PD and cholesterol in a POPE explicit membrane. **(H-J)** ACE2^+^ were pretreated with 10 mM of echinocystic acid, oleanolic acid, and ursolic acid for 1 h prior to transduction with pSARS-CoV-2 in the presence of each drug. After culture for 24 h, transduction efficiency was quantified by measuring the activity of firefly luciferase in cell lysates. Data are representative of three independent experiments with triplicate samples. Error bars indicate SEM. P values were determined by unpaired, two-tailed Student’s t-test. NS, not significant (H). Comparison of chemical structure of platycodin D with natural triterpene compounds without sugar moiety used in this experiment (I). ACE2^+^ were treated with 10 mM of each compounds as indicated for 24 h and then stained with filipin-III (J).

### PD inhibits exocytosis-mediated membrane fusion

As a proof-of-concept experiment, we tested the effect of PD on the well-known membrane fusion event of spontaneous inhibitory post-synaptic currents (sIPSC) in acute brain slices. We performed slice patch-clamp recordings from hippocampal CA1 pyramidal neurons (**Figure 5A**) and recorded sIPSCs before and during an application of PD (**Figure 5B**). We found that PD effectively inhibited the frequency of sIPSC without affecting the amplitude (**Figure 5C, D**). The time course of the inhibition by PD was relatively fast with about 10-15 second after the onset (**Figure 5C**). The concentration-response relationship showed that PD effectively inhibited sIPSC frequency with IC_50_ of 6.724 μM, whereas PD did not inhibit the amplitude of sIPSC (**Figure 5E, F**). These results support the idea that PD’s inhibitory action on pSARS-CoV-2-entry might be due to its ability to interfere with membrane fusion. What is the possible inter-molecular relationship between PD and cholesterol? Cholesterol appears to assist PD in incorporating into membrane as evidenced by a 2.5-fold increase in IC_50_ value for inhibition of pSARS-CoV-2-entry under lipoprotein-free condition (**Figure S4**). Taken together, these results strongly suggest that PD blocks pSARS-CoV-2-entry via interacting with cholesterol and preventing membrane fusion of viral membrane onto host-cell membrane.

**Figure 5.**
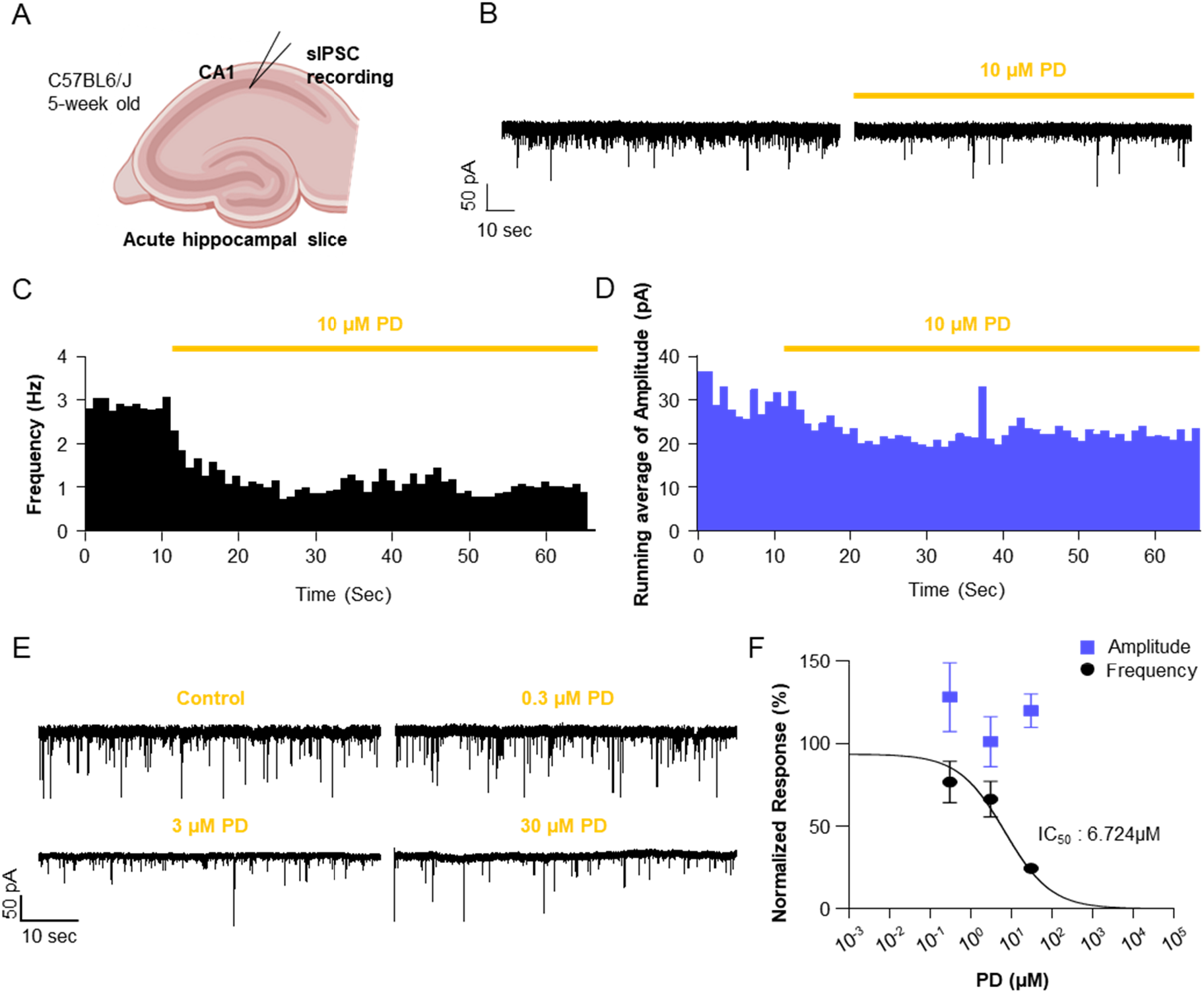
PD prevents membrane fusion as evidenced by a reduction of the frequency of spontaneous-inhibitory post-synaptic current (sIPSC) in the CA1 neurons. **(A)** Experimental scheme of sIPSC recording in CA1 hippocampus. **(B)** Representative traces of sIPSC recorded from CA1 hippocampus before 10 μM PD treatment (left) and during 10 μM PD bath application (right). **(C)** Running frequency during 1 h 10 μM PD treatment. sIPSC events were binned with one minute to represent as frequency in Hz. **(D)** Running average of amplitude during 1 h 10 μM PD treatment. sIPSC amplitudes were binned for every minute. **(E)** Representative traces of sIPSC recorded from CA1 hippocampus in control, 0.3 μM, 3 μM, 30 μM PD incubated slice, respectively. **(F)** Normalized responses for frequency (black) and amplitude (blue) of sIPSC in control (n=15), 0.3 μM (n=8), 3 μM (n=9), 30 μM (n=6) PD incubated conditions. Control condition data (0 μM) were not represented, but included in non-linear fitting.

### PD inhibits authentic SARS-CoV-2 infection into Vero and Calu-3 cells

As a final measure, we evaluated the antiviral activity of PD on the authentic SARS-CoV-2 viruses obtained from the Korea Centers for Disease Control and Prevention. We performed immunocytochemistry-based assessment of the infection of SARS-CoV-2, using an antibody against SARS-CoV-2 nucleocapsid N protein, as previously described (Ko et al., 2020). Infected host cells were automatically counted using an in-house image analysis software. For the host cells, we utilized both monkey-originated Vero and human-originated Calu-3 cell lines, whose western-blot analysis showed abundant expression of ACE2 in both cell types, the lack of TMPRSS2 expression in Vero cells, and high expression of TMPRSS2 in Calu-3 cells (**Figure 6A**). We found that PD caused a significant reduction in SARS-CoV-2 infection in both TMPRSS2-negative Vero and TMPRSS2-postive Calu-3 with IC_50_’s of 1.19 μM and 4.76 μM, respectively (**Figure 6B-D**). YGS also effectively inhibited SARS-CoV-2 infection into Vero cells with IC_50_ of 10.9 mg/ml (**Figure 6C, D**). These IC_50_ values obtained from the authentic SARS-CoV-2 were within the similar range as those IC_50_ values obtained from pSARS-CoV-2 (**Figure 1C, 2C**), indicating that PD is equally effective against the authentic SARS-CoV-2 viruses. Unlike other previously reported drugs (**Table 1**), PD shows the unique property of equal high potency against SARS-CoV-2 infection in both TMPRSS-negative Vero and TMPRSS-positive Calu-3 cells. This is in marked contrast to other drugs, which show only one-sided potency in either TMPRSS-negative Vero (chloroquine) or TMPRSS-positive Calu-3 (camostat, nafamostat). Taken together, these results indicate that PD has an important advantage over other known drugs in that it can potently prevent SARS-CoV-2 infection by inhibiting both lysosome- and TMPRSS2-driven SARS-CoV-2-entry pathways at the common process of viral membrane fusion by disrupting the host-cell membrane cholesterol (**Figure 7**).

**Figure 6.**
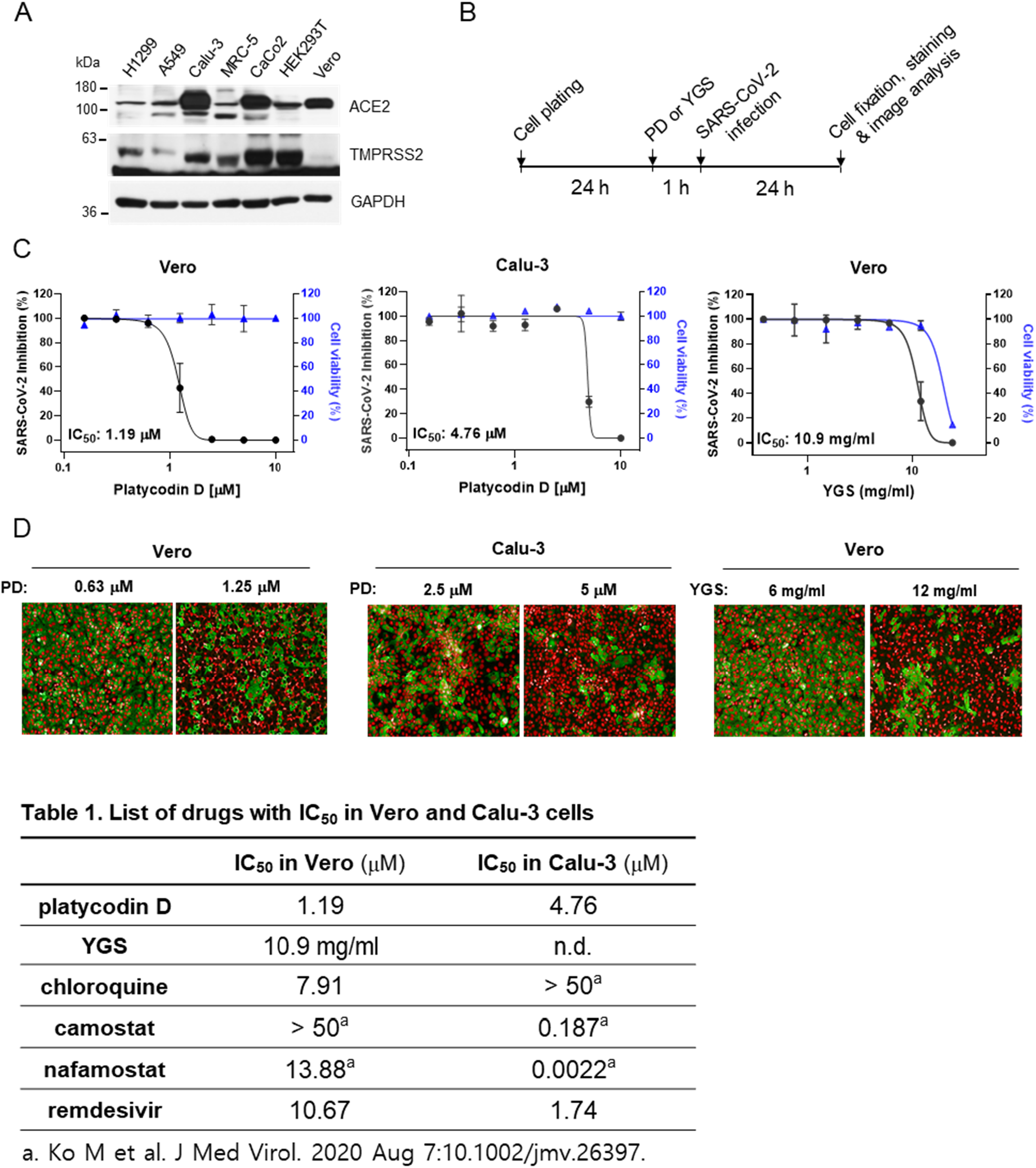
PD inhibits authentic SARS-CoV-2 infection into Vero and Calu-3 cells. **(A)** The protein levels of ACE2 and TMPRSS2 in multiple cell lines were assessed by western blot analysis. GAPDH was used as a loading control. **(B)** Experimental timeline for PD or YGS treatment and authentic SARS-CoV-2 infection before staining viral N protein and image analysis. **(C)** Dose-response curve analysis by immunofluorescence for PD and YGS. The black circles represent inhibition of authentic SARS-CoV-2 infection (%), and the blue triangles represent cell viability (%). Mean ± SEM was calculated from duplicate experiments. **(D)** The confocal microscope images show viral N protein (green) and cell nuclei (red) at concentrations near the IC_50_ of PD in Vero and Calu-3 cells, and near the IC_50_ of YGS in Vero cells.

**Figure 7.**
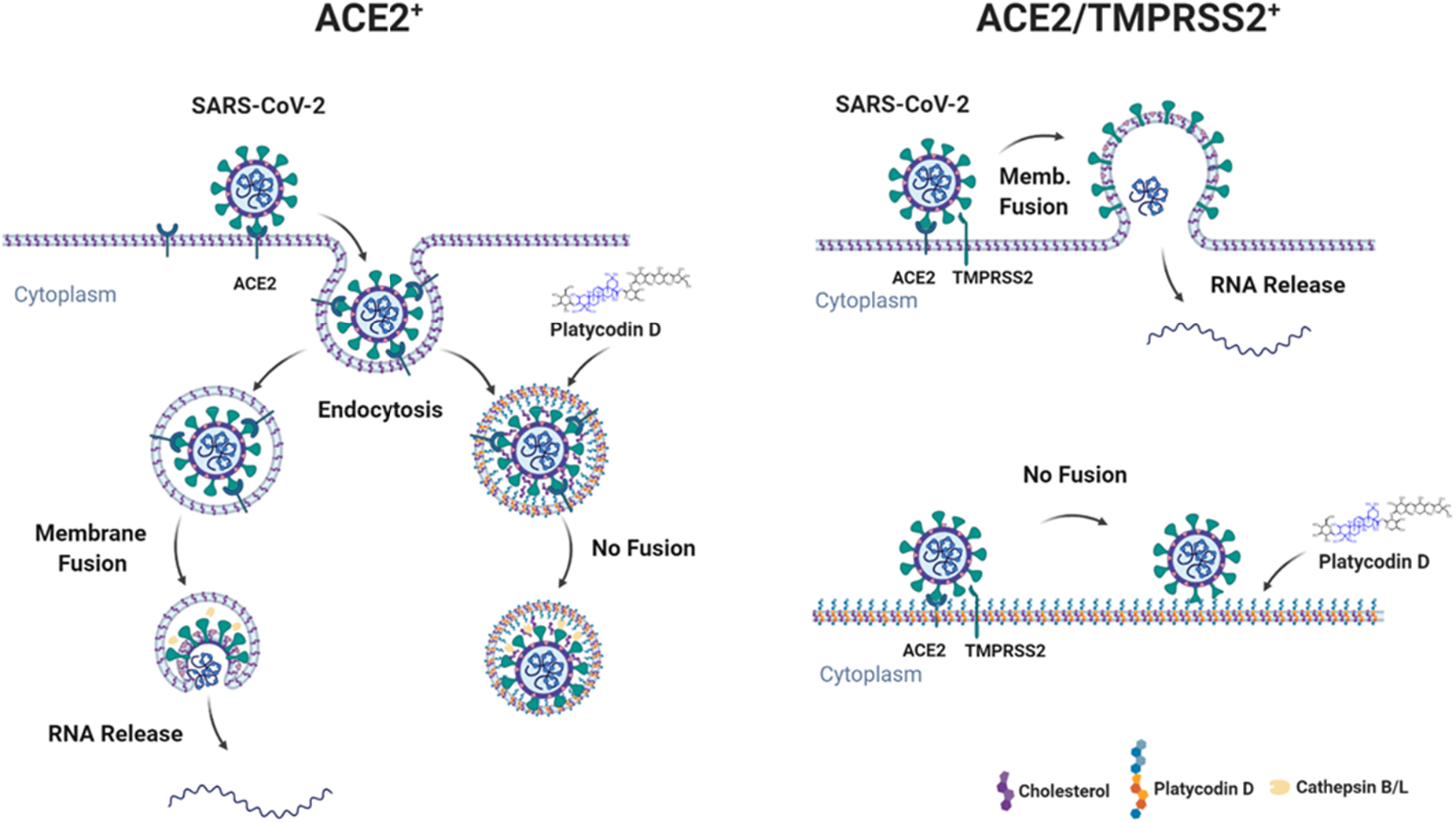
Schematic model of how PD prevents SARS-CoV-2-entry. **(A, B)** In both ACE2^+^ and ACE2/TMPRSS2^+^, PD incorporates into the host membrane with the sugar moieties protruding through the surface of membrane and inhibits SARS-CoV-2 membrane fusion with the endolysosomal membrane (A) and the plasma membrane (B).

## Discussion

SARS-CoV-2 binds to ACE2 and enters into host cells via membrane fusion after proteolytic cleavage of SARS-CoV-2 S-protein by 1) cathepsin B/L in the lysosomal membrane or 2) TMPRSS2 in the plasma membrane. We have demonstrated that PD effectively inhibits both of the SARS-CoV-2-entry pathways almost equally in ACE^+^ and ACE/TMPRSS2^+^ cells with IC_50_’s ranging from 0.69 μM to 4.76 μM against both pseudo- and authentic-SARS-CoV-2 viruses. To the best of our knowledge, PD is the first single compound that simultaneously blocks the two main pathways of SARS-CoV-2 infection. As the 1^st^ generation inhibitor, chloroquine was introduced as a potential inhibitor against SARS-CoV-2 infection. However, clinical trials with this drug have mostly failed because it blocks only cathepsin-mediated viral entry (Hoffmann et al., 2020b). To date, TMPRSS2 inhibitors such as camostat and nafamostat have been considered as the 2^nd^ generation inhibitor of SARS-CoV-2-entry. However, it has been recently reported that they are not effective in preventing viral entry into TMPRSS2-negative cells (Ko et al., 2020). Thus, we propose PD as the next generation inhibitor of SARS-CoV-2 infection, which blocks both of the entry pathways with much higher potency than the existing drug candidates such as chloroquine, camostat, and nafamostat.

One of the highlights of our study is that PD alters the distribution of membrane cholesterol, which contributes to PD’s anti-SARS-CoV-2 activity. Our finding is consistent with the very recent reports showing that 25-hydrocholesterol (25-HC) has antiviral effect on SARS-CoV-2 infection by accumulating cholesterol in the late endosomes and potentially restricting the S-protein-mediated membrane-fusion via depletion of membrane cholesterol (Wang et al., 2020b; Zang et al., 2020; Zu et al., 2020). These reports strongly support our notion that alteration of membrane cholesterol can be an effective strategy to prevent SARS-CoV-2 infection. However, it is still unclear how PD interacts with membrane cholesterol and inhibits membrane fusion. Although a pharmacological inhibition of NPC caused a depletion of membrane cholesterol (Figure 3I), PD did not deplete membrane cholesterol (Figure 3K). These results raise a possibility that PD’s action mechanism involves cholesterol-independent mechanism. On the other hand, PD’s potency decreases by 2.5 fold when cholesterol was depleted (Figure S4), suggesting that cholesterol might help PD as PD intercalates into membrane. Based on the molecular modeling, one can make a reasonable prediction that PD and cholesterol behave similarly because the hydrophobic triterpenoid saponin moiety of PD and cholesterol show similar size and hydrophobicity. The major structural difference comes from the fact that PD contains an additional elaborate sugar moiety that is strongly hydrophilic due to the multiple hydroxyl groups of the sugar moiety, which cholesterol lacks. This raises an interesting possibility that while PD behaves similarly as cholesterol within the lipid bilayer, PD behaves profoundly different outside the lipid bilayer, creating a physical hindrance due to the elaborate sugar tail that extends out from the membrane (Figure 4G). Considering the fact that the thickness of lipid bilayer is about 10 nm, the sugar tail of PD could extend out by about 2-3 nm (Figure 4G). Such conspicuous protrusions on the membrane could greatly hinder any membrane fusion events that require a close proximity and direct contact between two membrane structures, e.g., SARS-CoV-2 membrane and host-cell membrane. A classic example of such fusion event is the synaptic release due to a membrane fusion of synaptic vesicle membrane onto presynaptic terminal membrane during the exocytosis. Indeed, we found that PD can also inhibit the synaptic release event due to membrane fusion (Figure 5), supporting our hypothesis that PD’s sugar-tail-protrusion hinders membrane fusion. Future investigations are needed to systematically test this exciting possibility.

Recent studies have revealed that ACE2 and TMPRSS2 are highly expressed in nasal epithelial cells with lesser amounts in the lung tissues (Hou et al., 2020; Sungnak et al., 2020). These reports provide a plausible explanation for why viral loads of SARS-CoV-2 in the upper respiratory tract peaks before the onset of symptoms and why presymptomatic transmission occurs. Therefore, reducing the viral load in the upper respiratory tract of COVID-19 patients at the early and asymptomatic stages must be the most effective strategy to stop the current pandemic. Here, we tested an anti-SARS-CoV-2 infection activity of Yonggaksan (YGS), a commercially available herbal medicine containing PG as the main ingredient and obtained IC_50_’s of 3.17 and 3.72 mg/ml for ACE2^+^ and ACE2/TMPRSS2^+^ with pSARS-CoV-2, respectively, and 10.9 mg/ml in Vero cells with authentic SARS-CoV-2. The formulation of YGS is a fine powder and the traditional usage of YGS is to apply a spoonful of 300 mg YGS on the throat membrane without drinking water and wait for several minutes. Through this unique formulation and usage, an effective PD concentration can be reached at regions around throat mucus membrane and along the epithelium lining of upper respiratory airways. Alternatively, if applicable, PD can be easily and directly administered into the nose in the form of nasal drops or spray for a therapeutic purpose. These ideas should be tested immediately in animal models and COVID-19 patients. Therefore, we suggest that PG-containing herbal medicines or foods as well as PD compound itself can be a promising therapeutic option to exert a powerful brake on the spread of SARS-CoV-2 within and between individuals.

It has been predicted that the emergence of SARS-CoV-2 from the bat-originated SARS-CoV is probably due to naturally occurring mutations during its natural course of spread (Menachery et al., 2015). Such an alarming concept forebodes further emergence of more virulent versions of SARS-CoV-2. However, what is comforting from our study is that Mother Nature has already prepared natural products such as PD, which is capable of protecting human kind from SARS-CoV-2 infection. This raises an important point that it might be advantageous to find a cure from the vastly diverse natural products in addition to developing vaccines, which might become ineffective when new variants of SARS-CoV-2 emerges (Menachery et al., 2015).

## Acknowldegements

This research was supported by Institute for Basic Science (IBS), Center for Cognition and Sociality, Cognitive Glioscience Group (IBSR001-D2) and the National Research Foundation of Korea (NRF) grant funded by the Korean government (MSIT) (NRF-2017M3A9G6068245 and NRF-2020M3E9A1041756). The pathogen resource (NCCP43326) for this study was provided by the National Culture Collection for Pathogens. A patent application “Pharmaceutical composition for the prevention and treatment of COVID-19 comprising Platycodon grandiflorus root extracts” was filed with KIPO.

## Author contributions

Conceptualization: T.Y.K. and C.J.L.; Methodology: T.Y.K., C.J.L., S.K., S.H., and A.N.P.; Investigation: T.Y.K., Y.J., S.J., L.G., J.W., Y.H.J., S.K., M.W.J., W.W., and M.G.P.; Writing-original draft: T.Y.K. and C.J.L.; Writing-review and editing: T.Y.K., C.J.L., S.J., S.K., S.H., L.G., A.N.P., and J.W.; Supervision: C.J.L., S.K., S.H., and A.N.P.; Funding acquisition: C.J.L. and S.K.

## Supplemetary figure legends

**Figure S1.**
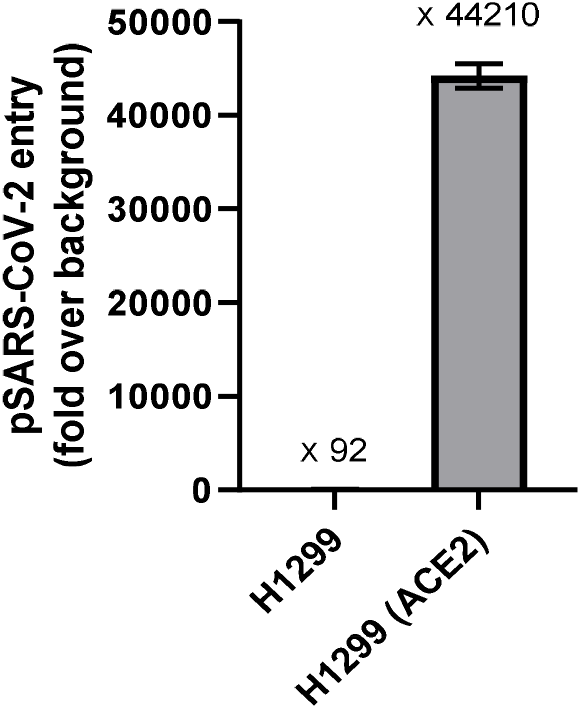
Transduction efficiency of pSARS-CoV-2. Parental H1299 cells and H1299 cells expressing ACE2 were transduced with pSARS-CoV-2 containing firefly luciferase gene. At 2-day post-transduction, transduction efficiency was determined by measuring luciferase activity in cell lysates. The activity obtained from untransduced cells were used for normalization.

**Figure S2.**
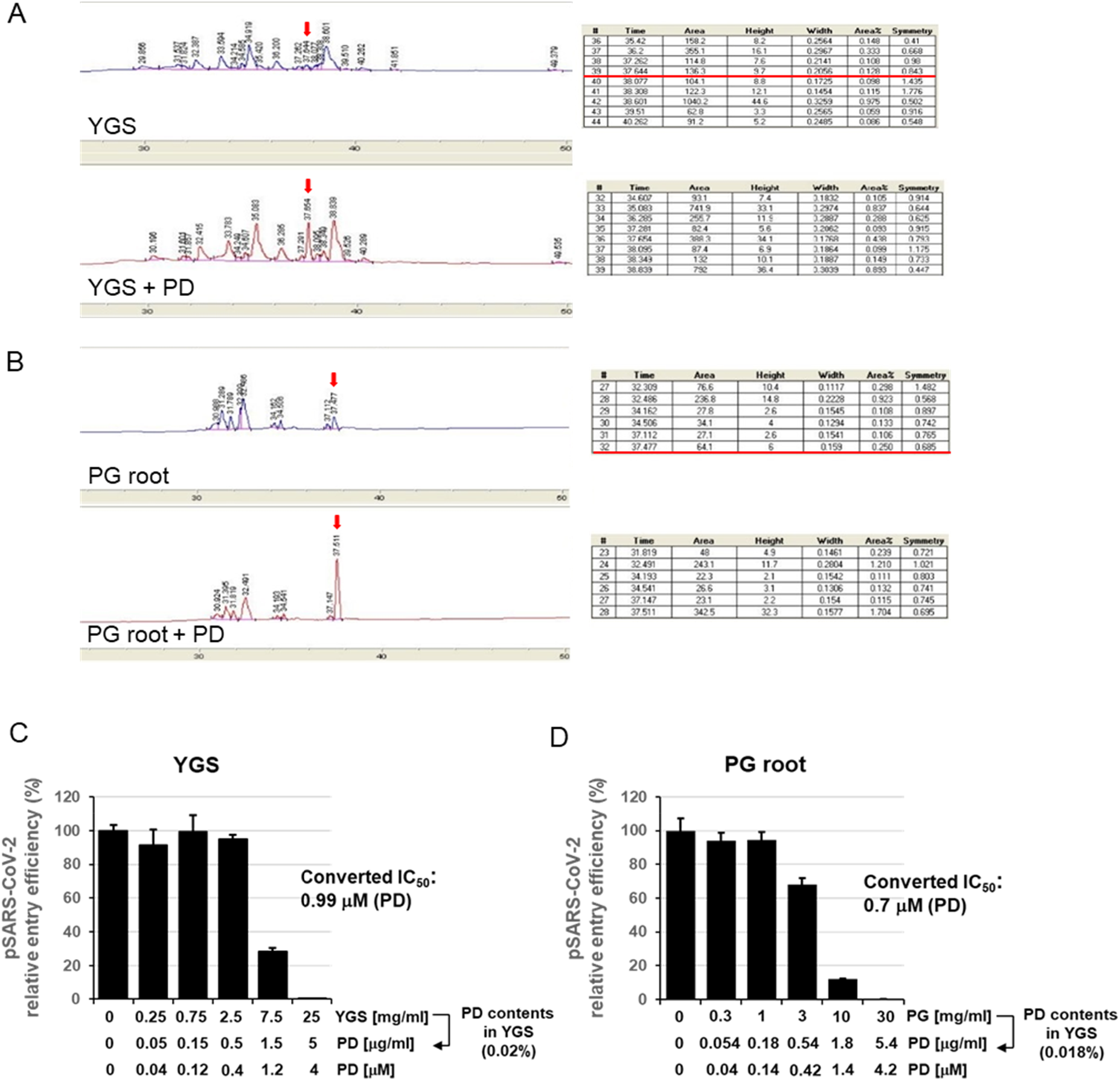
PD is the main compound that exhibits anti-SARS-CoV-2 activity in YGS and PG root. **(A)** Comparison of HPLC profiles for two samples of YGS (750 mg) and YGS (750mg) plus PD (0.25 mg) indicates that #39 (red arrow) is a peak for PD. The peak area for PD (red line in the table) is 136.3. **(B)** HPLC profiles of PG root (660 mg) and PG root (660 mg) plus PD (0.25 mg) indicates that #32 is a peak for PD and its value is 64.1. When these area value are put into the regression equation of “Y=2356*X-220.1” (R^2^ is 0.9989), which is obtained by a quantitative HPLC analysis using external standard calibration method (Figure S3), PD concentration can be calculated as 0.1513 mg/ml of PD in 750 mg of YGS (which is equivalent to 0.020%) and 0.1206 mg of PD in 660 mg of PG root (which is equivalent to 0.018%). **(C, D)** pSARS-CoV-2 entry assay using ACE2^+^ with serial three-fold dilutions of the stock solution of YGS and PG root. After converting mg/ml of YGS and PG root to mM of PD, the corrected IC_50_ of PD was 0.99 mM and 0.7 mM, respectively, which were very similar to IC_50_ of 0.69 mM obtained from the pSARS-CoV-2 entry assay with PD (Figure 1D), indicating that PD is the main compound that exhibits anti-SARS-CoV-2 activity in the YGS powder and PG root.

**Figure S3.**
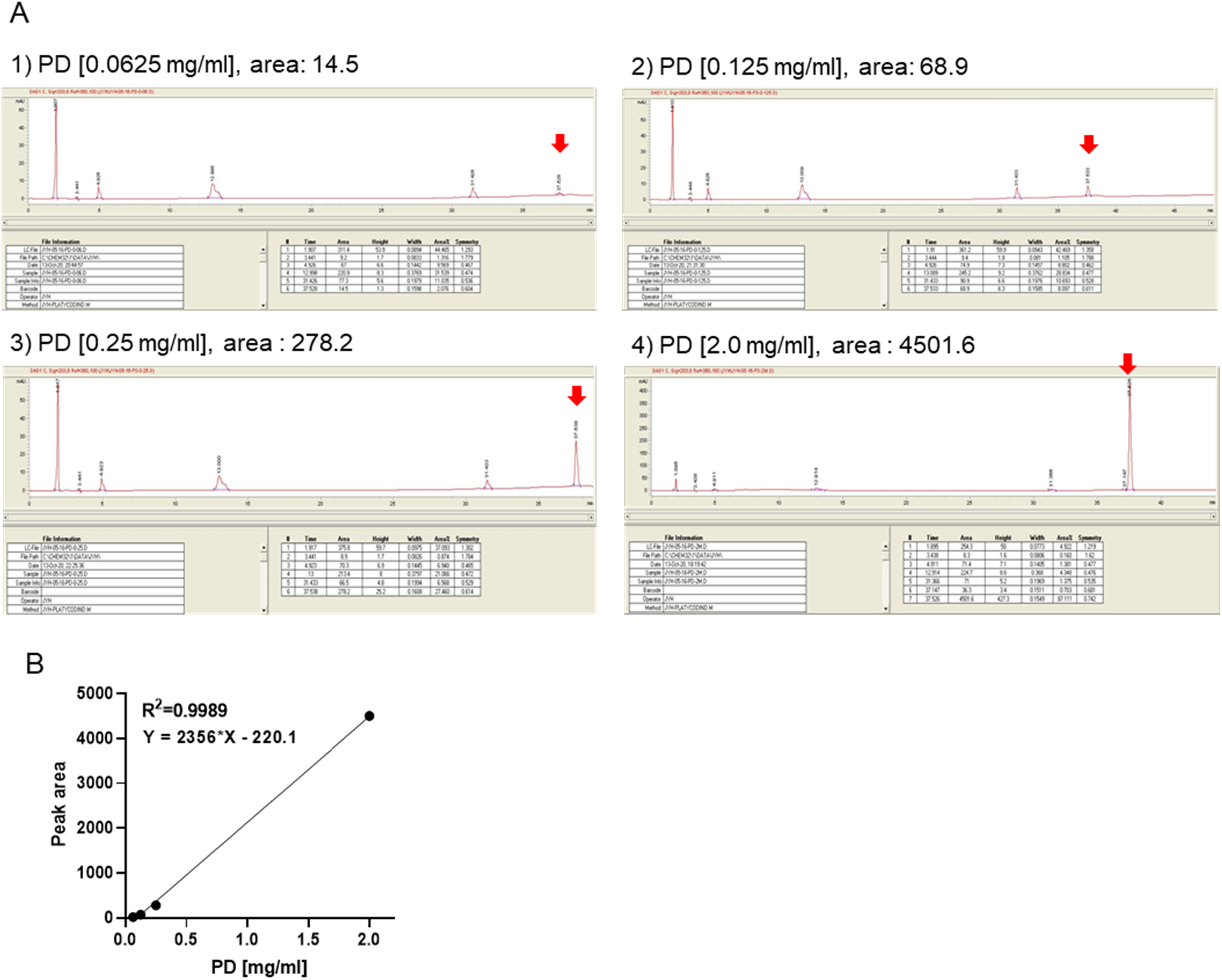
Quantification of PD by HPLC using external standard calibration method. **(A, B)** HPLC quantitative analysis using external standard method. Calibration curve (B) was created with values of the peak area for 0.0625, 0.125, 0.25, and 2.0 mg/ml of PD (A). R2 value and regression equation was obtained as 0.9989 and “Y=2356*X-220.1”, respectively (B).

**Figure S4.**
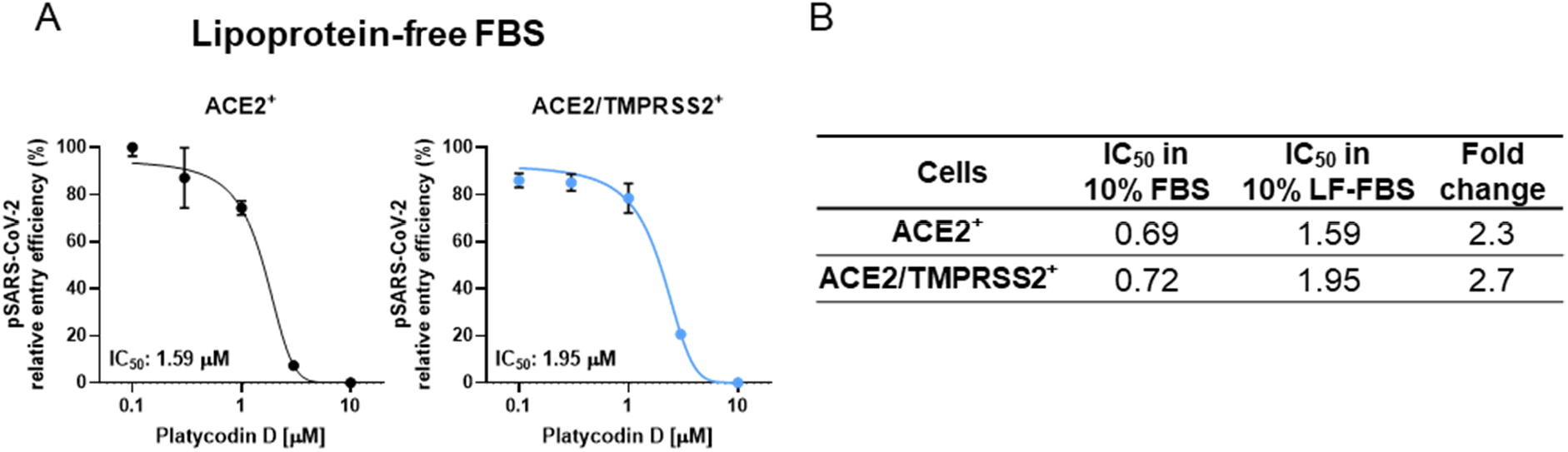
PD’s inhibitory action against SARS-CoV-2-entry is cholesterol-dependent. **(A)** ACE2^+^ and ACE2/TMPRSS2^+^ cultured in 10% lipoprotein-fee FBS (LF-FBS)-containing media for 24 h were pre-treated with indicated concentrations of PD for 1 h and further transduced with pSARS-CoV-2 in 10% LF-FBS-containing media in the presence of PD. After culture for 24 h, transduction efficiency was quantified by measuring the activity of firefly luciferase in cell lysates. Data are representative of two independent experiments with triplicate samples. Error bars indicate SEM. **(B)** Comparison of IC_50_ of PD between in normal culture condition (Fig. 1d, l) and in lipoprotein-free culture condition.

## Materials and Methods

### Cell culture

H1299, A549, MRC-5, and CaCo2 cells were obtained Korean Cell Line Bank. HEK293T cells were obtained from the American Type Culture Collection (ATCC, USA, CRL-3216). H1299 and A549 cells were cultured in RPMI-1640 (Gibco, USA) and MRC-5, CaCo2, and HEK293T cells were cultured in Dulbecco’s Modified Eagle’s Medium (DMEM; Corning, USA), supplemented with 10% fetal bovine serum (FBS) (Gibco, USA) and 1X Penicillin-Streptomycin solution (HyClone, USA) at 37°C in a humidified incubator containing 5% CO2. For cell culture in lipoprotein-free condition, lipoprotein depleted FBS (Kalen biomedical, USA) was used. For authentic SARS-CoV-2 experiments, Vero cells were obtained from ATCC (CCL-81) and maintained at 37°C with 5% CO2 in DMEM supplemented with FBS and 1X Antibiotic-Antimycotic solution (Gibco, USA). Calu-3 used in this study is a clonal isolate, which shows higher growth rate compared to the parental Calu-3 obtained from ATCC (HTB-55). Calu-3 was maintained at 37°C with 5% CO2 in Eagle’s Minimum Essential Medium (EMEM, ATCC), supplemented with 10% FBS and 1X Antibiotic-Antimycotic solution

### Reagent

Platycodin D, U18666A, E64d, chloroquine, camostat mesylate, nafamostat mesylate, ginsenoside Rb1, ginsenoside Rb3, ginsenoside mix, glycyrrhizin, isoliquiritigenin, echinocystic acid, oleanolic acid, ursolic acid, and methyl-beta-cyclodextrin were purchased from Sigma-Aldrich Co (USA).

### Plasmids and establishment of stable cell line

The full length of ACE2 sequence from pCEP4-myc-ACE2 (a gift from Erik Procko, Addgene plasmid #141185) was cloned into pHR-CMV lentiviral expression vector (a gift from A. Radu Aricescu, Addgene plasmid # 113888) via EcoRI and AgeI restriction sites. TMPRSS2 lentiviral expression vector, RRL.sin.cPPT.SFFV/TMPRSS2 (variant 1).IRES-neo.WPRE (MT130) was gifted by Caroline Goujon (Addgene plasmid # 145843). For gene silencing, the oligonucleotides that contain shRNA sequence targeting NPC1 (5’-CCAGGTTCTTGACTTACAA-3’), NPC2 (5’-CGGTTCTGTGGATGGAGTTAT-3’) and nonspecific (NS) (5’-CAACAAGATGAAGAGCACCAA-3’) was cloned into pLKO.1 puro lentiviral shRNA plasmid. For stable cell line generation, these plasmids were transiently transfected into HEK293T cells with packaging plasmid psPAX2 and envelope plasmid pMD2.G using Lipofectamine 3000 transfection reagent (Thermo Fisher, USA) according to the manufacturer’s instructions. At 24 h and 48 h post-transfection, the lentivirus particles containing supernatants were harvested, filtered through 0.45 μm-pore-size filters and were used to infect H1299 cells which were seeded on 6 well plates and cultured until reaching 70-80% confluence. One milliliter of the virus supernatants was directly overlaid on the cells in the presence of polybrene (Merck, Germany) at a final concentration of 4 μg/ml. After 24 h, the supernatants were changed with fresh medium and cultured for 2~3 days.

### SARS-CoV-2 Spike-pseudotyped lentivirus production and transduction

SARS-CoV-2 Spike (S)-pseudotyped lentiviruses was generated using 2th generation lentiviral packing system. In brief, HEK293T cells that reached 70-80% confluency in a 6-well plate were transfected with 1 μg of lentiviral backbone that contains expression cassettes for firefly luciferase, 0.75 μg of psPAX2 packing plasmid, and 0.5 μg of SARS-CoV-2 Spike plasmid (a gift from Fang Li, Addgene plasmid #145032) using Lipofectamine 3000 transfection reagent (Invitrogen, USA) following the manufacturer’s instructions. pMD2.G plasmid was used to create control lentivirus pseudotyped with VSV-G. At 24 h and 48 h post-transfection, supernatants containing SARS-CoV-2 Spike-pseudotyped virus particles were collected, filtered through a 0.45 um pore size filter, and stored at 4°C until use. For transduction, H1299 cells stably expressing ACE2 or ACE2 plus TMPRSS2 were cultured in 48 well plate until reaching 70-80% confluence. The virus supernatants were directly overlaid on H1299 cells in the presence of polybrene (Merck, Germany) at a final concentration of 4 μg/ml. For experiments with drugs, H1299 cells were pre-treated for 1 h with each drugs before transduction. After 24 h incubation, transduction efficiency was quantified by measuring the activity of firefly luciferase in cell lysates using a luciferase assay system (Promega, USA) and SpectraMax iD5 Multi-Mode Microplate Reader (Molecular Devices, USA).

### Authentic SARS-CoV-2 virus and dose-response curve analysis by immunofluorescence assay

The Korean strain of SARS-CoV-2 (βCoV/KOR/KCDC03/2020) was provided from Korea Centers for Disease Control and Prevention (KCDC), and was propagated in Vero cells. Viral titers were determined by plaque assays in Vero cells. All experiments using SARS-CoV-2 were performed at the Institut Pasteur Korea in compliance with the guidelines of the KNIH, using enhanced Biosafety Level 3 (BSL-3) containment procedures in laboratories approved for use by the KCDC. Vero cells were seeded at 1.2 × 10^4^ cells per well and Calu-3 cells were seeded at 2.0 × 10^4^ cells per well in black, 384-well, μClear plates (Greiner Bio-One, Austria), 24 h prior to the experiment. Ten point DRCs were generated, with compound concentrations ranging from 0.02–10 μM and 0.05-24 mg/ml. Except for chloroquine and remdesivir which ranges from 0.1-150 μM. For viral infection, plates were transferred into the BSL-3 containment facility and SARS-CoV-2 was added at a multiplicity of infection (MOI) of 0.0125 in Vero cells and 0.5 in Calu-3 cells. The cells were fixed at 24 hpi with 4% paraformaldehyde (PFA), stained with SARS-CoV-2 nucleocapsid (N) protein, and analyzed by immunofluorescence analyses. The acquired images were analyzed using Columbus software (PerkinElmer, Inc. Waltham, MA) to quantify cell numbers and infection ratios, and antiviral activity was normalized to positive (mock) and negative (0.5% DMSO) controls in each assay plate. DRCs were fitted by sigmoidal dose-response models, with the following equation: Y = Bottom + (Top - Bottom)/(1 + (IC_50_/X)Hillslope), using XLfit 4 Software or Prism6. Half-maximal inhibitory concentration (IC_50_) values were calculated from the normalized activity dataset-fitted curves. IC_50_ were measured in duplicate, and the quality of each assay was controlled by Z’-factor and the coefficient of variation in percent (%CV).

### Cell viability assay

H1299, Calu-3, Vero cells seeded in 96-well plate (5×10^3^ cells/well) were treated with the indicated concentrations of PD. After 24 h treatments, WST-8 solution (Biomax, Korea) was added and incubated for 2 h. Water-soluble formazan formed in the culture medium was measured by SpectraMax iD5 Multi-Mode Microplate Reader (Molecular Devices, USA) at 450 nm absorbance. The relative cell viability (%) was expressed as a percentage relative to the DMSO-treated control cells.

### Protein extraction and Immunoblot Analysis

Cell lysates were prepared by solubilizing cells with ice-cold RIPA lysis buffer (Rockland Immunochemicals, USA) supplemented with protease and phosphatase inhibitor cocktail (Thermo Fisher Scientific, USA), followed by centrifugation at 13,000 rpm for 20 min. The cleared cell lysates were mixed with a proper volume of 5x SDS sample buffer, separated by SDS-PAGE, and transferred to nitrocellulose membranes. After blocking with 5% skim milk in TBST (20 mM Tris-HCl (pH 7.5), 150 mM NaCl, 0.05% Tween 20) for 1 h at room temperature (RT), the membranes were incubated in the in TBST at 4°C overnight with the following primary antibodies: rabbit anti-NPC1 (Novus, NB400-148), rabbit anti-NPC2 (Novus, NBP1-84012), rabbit anti-ACE2 (Abcam, ab15348), rabbit anti-TMPRSS2 (Abcam, ab92323), and mouse anti-GAPDH (Abcam, ab8245). The corresponding horseradish peroxidase-conjugated secondary antibodies (KPL, USA) were incubated 1 h at RT. Antibody-protein complexes were detected using ECL Western Blotting Substrate (Thermo Fisher Scientific, USA).

### Filipin III cholesterol staining

H1299 cells grown on coverslips were fixed with 4% PFA at RT for 10 min and then stained with 5 μg/ml filipin-III (Cayman, USA) in PBS/1% FBS solution at RT for 2 h in the dark. The stained cells were examined under LSM700 confocal microscope (Carl Zeiss, Germany).

### Flow cytometry analysis

Expi293F cells were transfected with pcDNA3-SARS-CoV-2-S-RBD-sfGFP (a gift from Erik Procko, Addgene plasmid # 141184) using ExpiFectamine™ 293 Transfection Kit according to manufacturer’s directions (Thermo Fisher Scientific, USA). The cells were then cultured for 4-5 days and removed by centrifugation at 800×g for 5 minutes, and the culture medium was stored at −4 °C. To analyze the effects of PD on binding of CoV-2-RBD-GFP to cell surface ACE2, H1299 cells expressing ACE2 were treated with DMSO or 5 μM PD for 1 h and washed with ice-cold PBS-BSA, followed by incubation with a 1/10 dilution of medium containing CoV-2-RBD-GFP for 30 minutes on ice. The H1299 cells were then washed twice with PBS-BSA and analyzed on a BD LSRFortessa™ Flow Cytometer.

### HPLC analysis

The root of Platycodon grandiflorum or YGS powder was dissolved in distilled water and analyzed using an HPLC system (Agilent 1100) at 203 nm with a RS Tech HECTOR-M C18 column (4.6 x 250 mm, 5 micron particle size, RS Tech Corp, Cheongju, South Korea). The column was eluted at 25 °C with a mixture of solvent A (water) and solvent B (acetonitrile). The gradient eluent system consisted of 82:18 (A:B) from 0 to 22 min, 82:18 (A:B) to 70:30 (A:B) from 22 min to 32 min, 70:30 (A:B) to 50:50 (A:B) from 32 min to 60 min at a flow rate of 1 mL/min.

### Molecular modelling of MβCD inclusion complexes

Schrodinger Maestro software 2017 suite (Maestro, Schrödinger, LLC, New York, NY) was used to prepare and score the MβCD binding complexes. The software tools were applied using the default settings and pH 7.4, unless stated otherwise below. The binding complex models were prepared based on the crystal structure of the β-cyclodextrin (βCD) cholesterol inclusion complex (CSD entry: KEXQUC) (Christoforides et al., 2018). The crystal structure of tetradeca-2,6-O-methyl-β-cyclodextrin (CSD entry: BOYFOK03) (Stezowski et al., 2001) was used to represent MβCD in the binding complex. Both crystal structures were obtained from the Cambridge Structural Database (CSD) and all water molecules were removed. The ligands cholesterol and PD were drawn in ChemDraw Professional 16.0 and imported as structure data files into the Maestro LigPrep module. The Ligprep module was used to prepare all ligands for further use: determine the ligand partial charges, optimize the geometry and perform energy minimization using the OPLS3 force field. The MβCD binding complexes were prepared using the βCD-cholesterol complex as a template. First, the atomic coordinates of the two template βCD units were copied onto two corresponding MβCD units. Cholesterol was added inside the complex by copying atomic coordinates of the template guest cholesterol molecule, whereas PD was placed using a rigid alignment, maximum common structure molecular overlay tool. To account for the increase of the macrocycle size, as well as remove molecular strain and atomic bumps and relax the binding complex structures, a restrained energy minimization algorithm was applied using the OPLS3 force field and heavy atom RMSD restriction of 2 Å. The obtained ligand binding poses were scored in place by the Glide SP (standard precision) dock score and afterwards the relative ligand binding affinity was estimated by the Prime MM-GBSA tool with a variable-dielectric generalized Born (VDGB) continuum solvation model and OPLS3 force field. The binding affinity is calculated using the equation: ΔG_bind_ = E_complex(minimized)_-(E_ligand(minimized)_ + E_host(minimized)_). More negative scores indicate stronger binding. The binding complex visualization was done using the Discovery Studio Client 2020 package (Dassault Systèmes; BIOVIA. Discovery Studio Modeling Environment; Release 2020; Dassault Systèmes: San Diego, CA, 2020).

### PD and cholesterol orientation in the cell membrane

The Discovery Studio Client 2020 package (Dassault Systèmes; BIOVIA. Discovery Studio Modeling Environment; Release 2020; Dassault Systèmes: San Diego, CA, 2020) was used to optimize the position and orientation of PD and cholesterol in an explicit membrane bilayer, consisting of 1-palmitoyl-2-oleoyl-sn-glycero-3-phosphoethanolamine (POPE) lipid molecules. Before running the membrane addition and molecule orientation algorithm, the 3D conformations of PD and cholesterol were prepared using the standard ligand preparation protocol and energy minimized with the general CHARMm force field. The molecules were first re-oriented to an implicit membrane by doing a step-wise search for the minimum solvation energy of the molecule, which was calculated by the charmm36 force field and Generalized Born Implicit Membrane (GBIM) module. The membrane thickness was set to 35 Å. After finding the optimal molecule position in the implicit membrane, the lipid bilayer was constructed by adding the POPE lipid molecules, water molecules and counterions (Na^+^ and Cl^-^) without performing system equilibration.

### Electrophysiology

For the brain acute slicing, 5 weeks C57BL/6J mice were anesthetized with isoflurane and decapitated. The The 300 μm transverse slice of hippocampus were prepared in ice-cooling NMDG-based cutting solutions (93 mM NMDG; 2.5 mM KCl; 1.2 mM NaH2PO4; 30 mM NaHCO3; 20 mM HEPES; 25 mM Glucose; 5 mM Sodium ascorbate, 2 mM Thourea, 3 mM Sodium pyruvate, 10 mM MgCl2, 0.5 mM CaCl2, pH adjust to 7.3 with HCl, 310 mOsm) on D.S.K Linear Slicer pro7 (Dosaka EM Co., Ltd) and recovered 15 minutes at 32°C in the same solution. Slices were re-recovered in oxygenated ACFS solution (126 mM NaCl; 24 mM NaHCO3; 2.5 mM KCl; 1 mM NaH2PO4; 2 mM MgCl2; 10 mM Glucose). Spontaneous IPSC was recorded under oxygenated ACFS solution by whole-cell voltage-clamp. Recording electrodes (6-8 MΩ) fabricated from standard-wall borosilicate glass (GC150F-10, Warner Instrument Corp., USA) were filled with an CsCl-based internal solution (135 mM CsCl; 4 mM NaCl; 0.5 mM CaCl2; 10 mM HEPES; 5 mM EGTA, 0.5 mM Na2-GTP; 2 mM Mg-ATP; 1 mM QX-314). For the acute PD treatment and sIPSC recording, 10uM PD containing-ACSF were applicated after setup the baseline. For the long-term PD treatment and sIPSC recording, slices were incubated in PD containing-ACSF (0, 0.3, 1, 3, 10, 30 μM, respectively) at least c one hour and whole-cell voltage-clamp recording in the same solutions. sIPSC was recorded at least 5 minutes. For sIPSC frequency and amplitude analyzing, Mini Analysis Program software (Synaptosoft) were used. Under −300 pA of holding current at baseline data were rule out for analysis.

### Statistical analysis

Data in this study were representative of two or three independent experiments with triplicate samples and presented as the mean± SEM. Statistical analysis was performed using student’s t-test or one-way ANOVA followed by Tukey’s post hoc test. A p-value less than 0.05 was considered statistically significant (*P < 0.05; **P < 0.01; ***P < 0.001; ****P < 0.0001; NS, not significant). For all statistical analyses, the Prism v.9.0.0 was used (GraphPad Software).

